## Supplementary information for "Spectro-temporal acoustical markers differentiate speech from song across cultures"

**The PDF file includes:**

Figs. S1 to S12

Statistics for Fig. 3

Tables S1 and S2

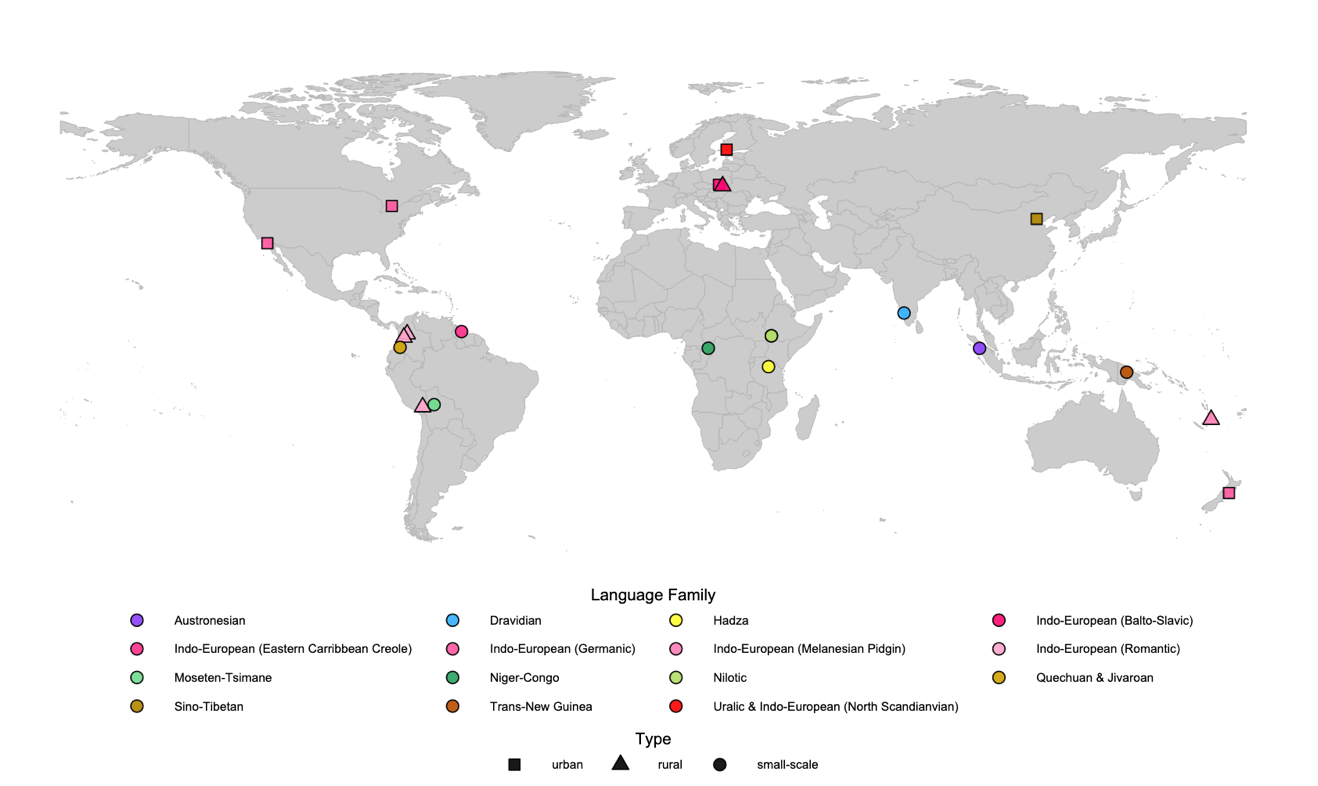

**Fig. S1: Societies from which vocalizations were recorded**. Triangles denote rural societies; circles denote small scale societies and squares denote urban societies, from ^6^, used with permission.

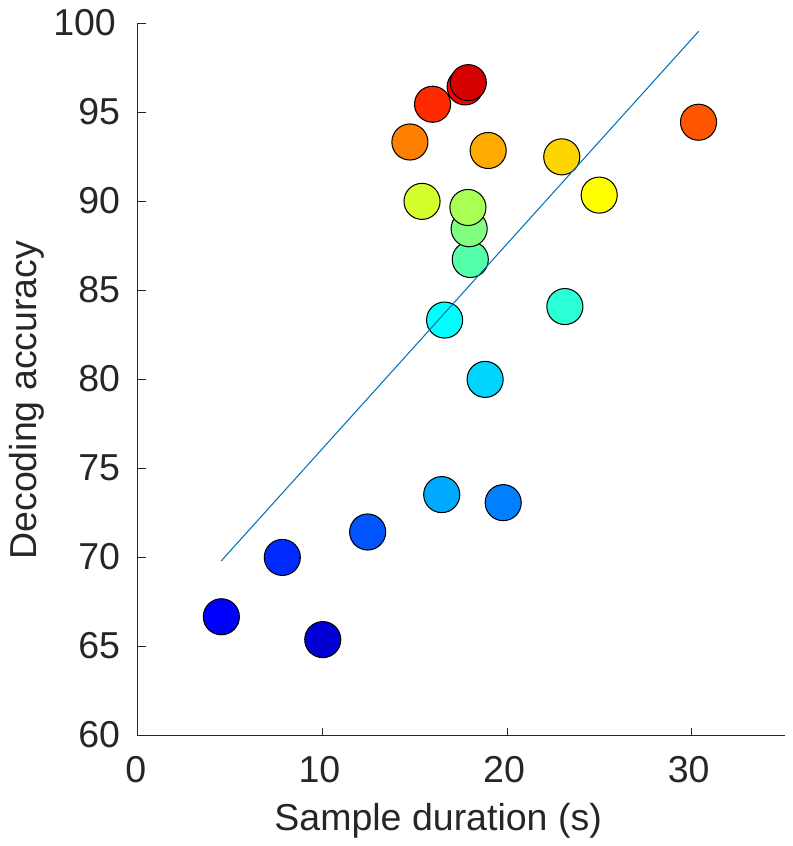

**Fig. S2:** Scatter plot of SVM decoding accuracy (Fig. 2D) against average sample duration (s). Colored circles represent each of the 21 societies/cultures (sorted as a function of the SVM decoding accuracy of Fig. 2D)

Statistics for Fig. 3:

*i)* Countries (Wilcoxon rank test, two-tailed, W(17) = 171 p <.001; Rank biserial correlation = 1.00; 95% Confidence Interval = [30.1 40.3] - Fig. 3A; accuracy = 84.7% ± 9.37 (SD); sensitivity = 83.5% ± 16.2 , specificity = 85.9% ± 12.5

*ii)* Language Families (Wilcoxon rank test, two-tailed, W(14) = 120, p <.001; Rank biserial correlation = 1.00; 95% Confidence Interval = [32.0 42.5]- Fig. 3B; accuracy = 87% ± 8.4 (SD); sensitivity = 87.2% ± 13.6, specificity = 86.8% ± 13.1

*iii)* World subregions (Wilcoxon rank test, two-tailed, W(13) = 105, p <.001; Rank biserial correlation = 1.00; 95% Confidence Interval = [26.5 38.3] - Fig. 3C; accuracy = 82.2% ± 9.56 (SD); sensitivity = 81.3% ± 16.5, specificity = 83.1% ± 13

*iv)* World regions (Wilcoxon rank test, two-tailed, W(5) = 21, p =.031; Rank biserial correlation = 1.00; 95% Confidence Interval = [28.3 39.9]; - Fig. 3D; accuracy = 86.2% ± 4.3 (SD); sensitivity = 84.2% ± 12.8, specificity = 88.2% ± 6.

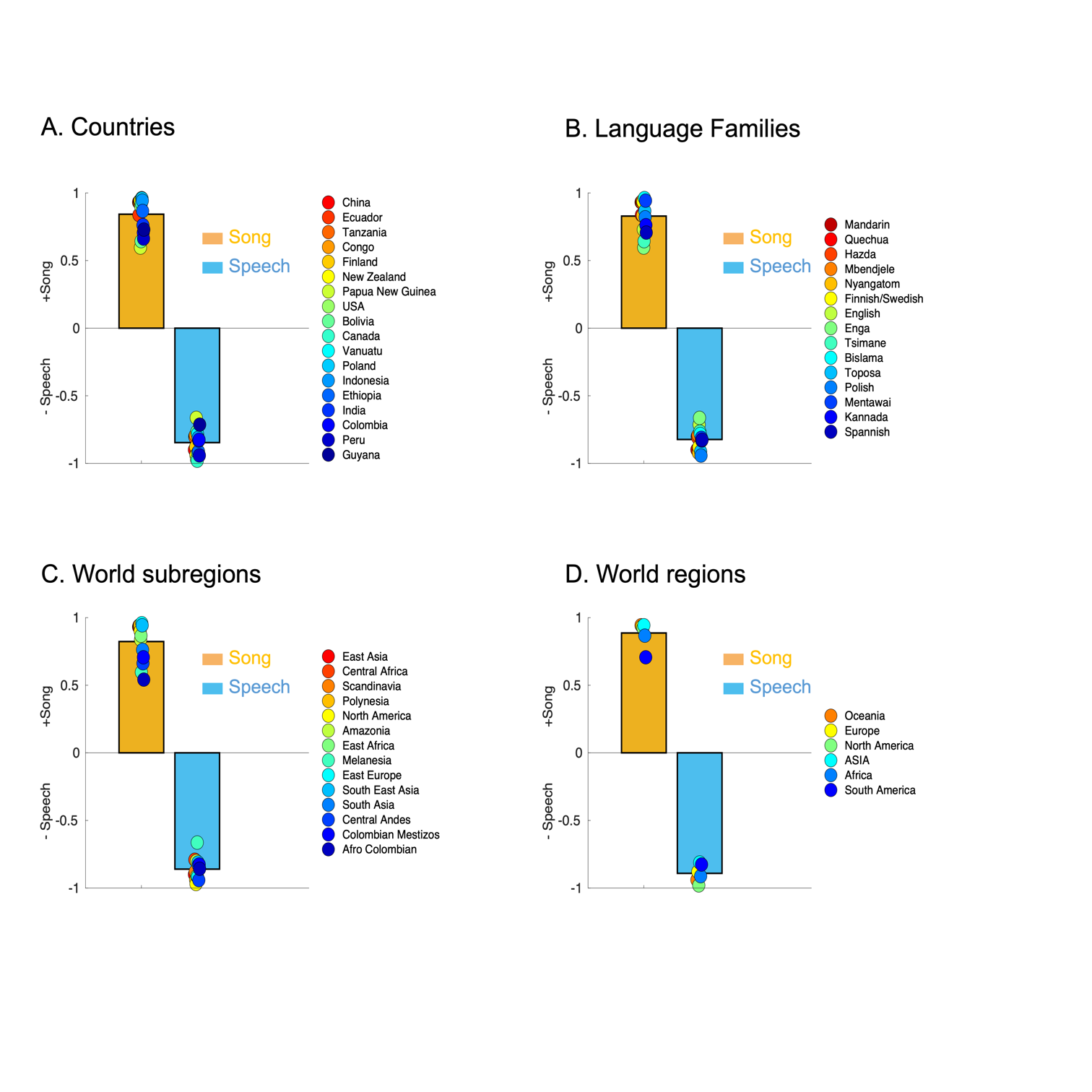

**Fig. S3: Behavioral ratings.** A. Countries-level behavioral ratings (chance level – 0) for song (orange) and speech (blue) samples. Colored circles represent the behavioral ratings for each country (sorted as a function of decoding accuracy in Fig. 3). B. C. D. same as (A.) for language families, world subregions, and world regions respectively.

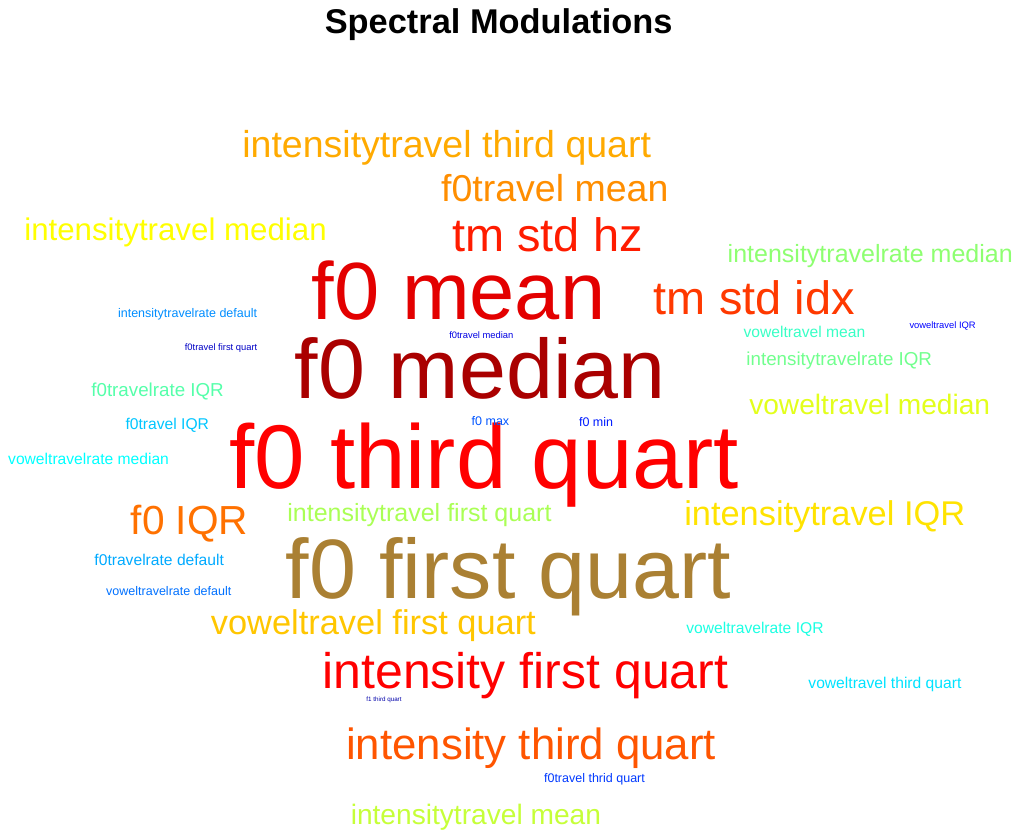

**Fig. S4. Word cloud presenting variables with the largest influence in the PLS** (before grouping them by label – see Table S2) for the spectral peak (S) in Fig. 5A. Font size varies as a function of VIP scores.

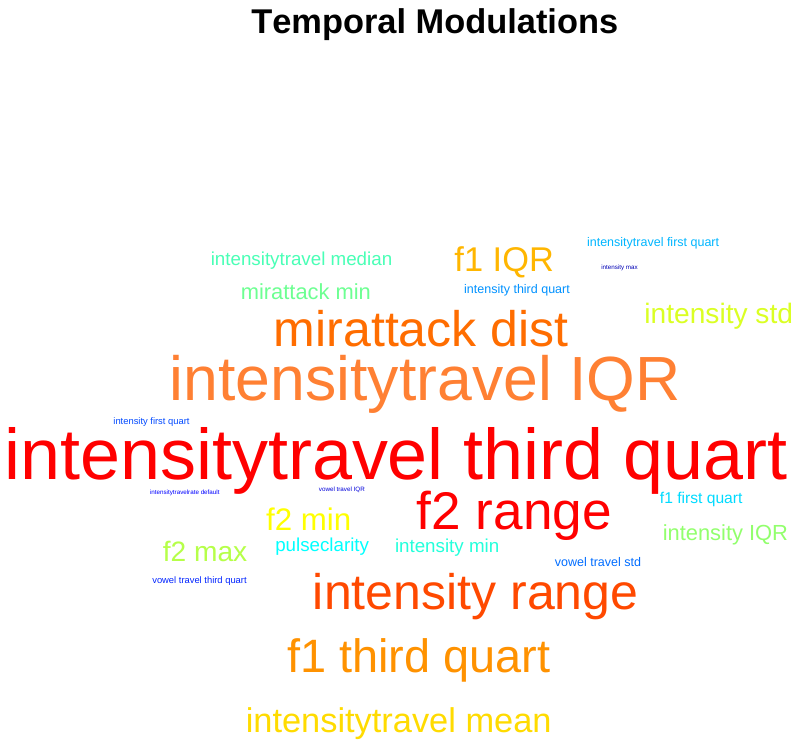

**Fig. S5. Word cloud presenting variables with the largest influence in the PLS** (before grouping them by label – see Table S2) for the temporal peak (T) in Fig.5 A. Font size varies as a function of VIP scores

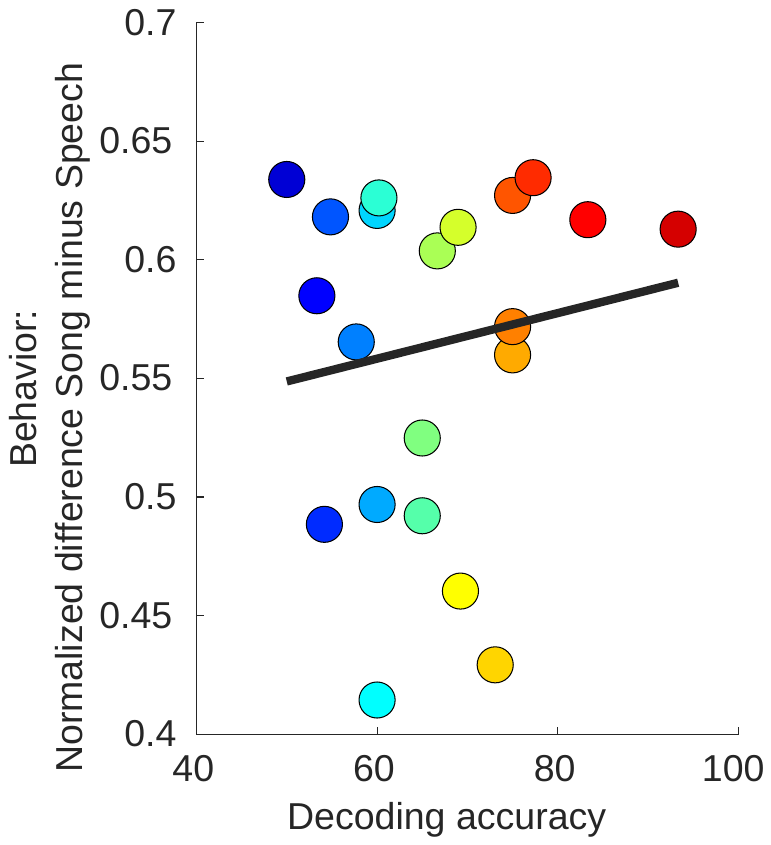

**Fig. S6:** **SVM decoding accuracy with acoustical variables as features does not predict behavior.** Scatter plot of SVM decoding accuracy (Fig. 5D, acoustical features only) against behavioral normalized difference (Song vs. Speech). Colored circles represent each of the 21 societies/cultures (sorted as a function of the SVM decoding accuracy of Fig. 2D)

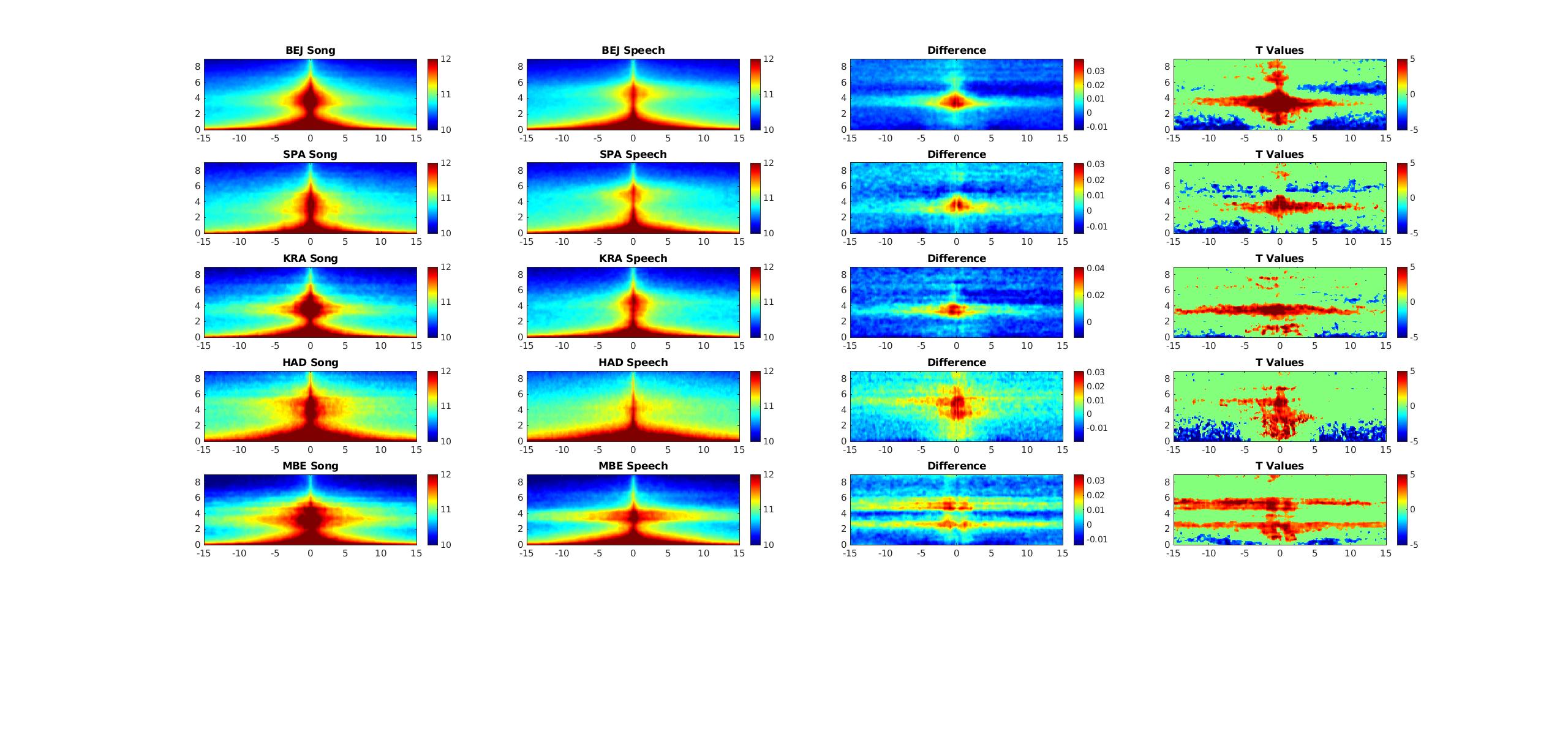

**Fig. S7. Extraction of spectro-temporal modulations patterns for singing and speaking vocalization samples.** For each sample (n indicates the number of different speakers for each society) we extracted the spectrotemporal modulation patterns and then averaged them. We computed the normalized difference between the song and speech modulation patterns and also performed non-parametric permutation statistics (uncorrected) for illustration. The panels in the third and fourth columns therefore show where in the modulation space song differs from speech (red part of the scale) and where speech differs from song (blue part of the scale). Societies are sorted as a function of decoding accuracy presented in Fig. 2.

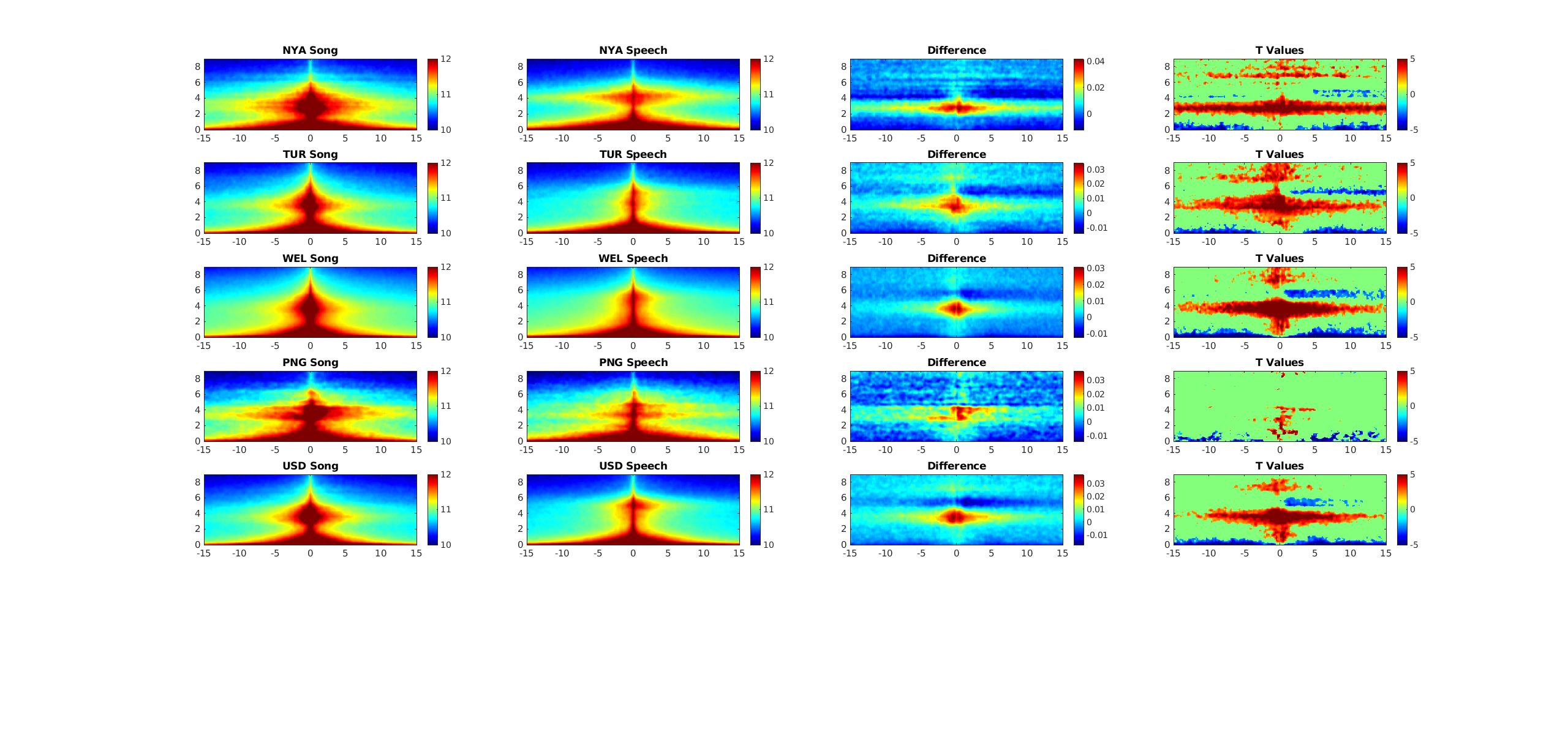

**Fig. S8. Extraction of spectro-temporal modulations patterns for singing and speaking vocalization samples.** Same as Fig. S7

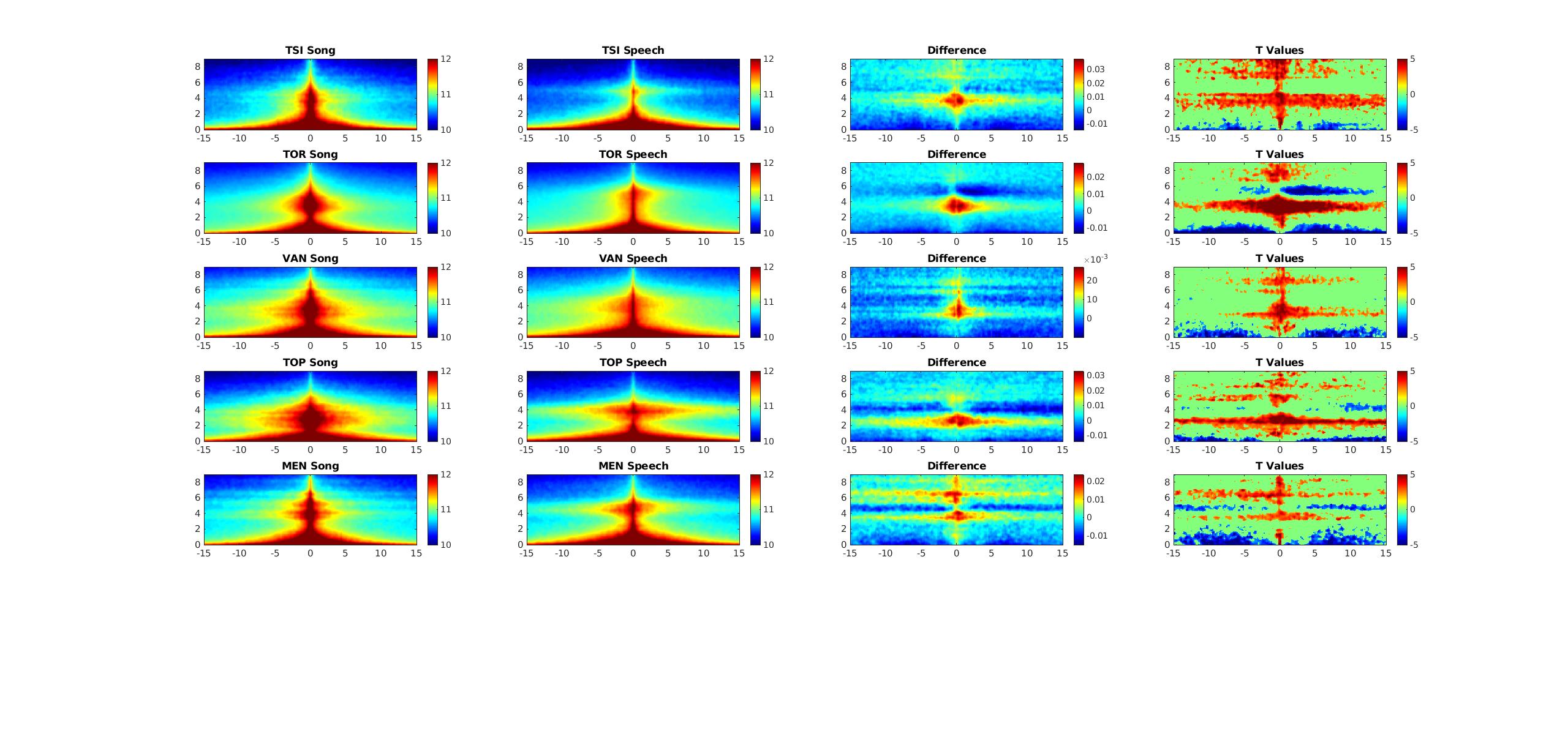

**Fig. S9. Extraction of spectro-temporal modulations patterns for singing and speaking vocalization samples.** Same as Fig. S7

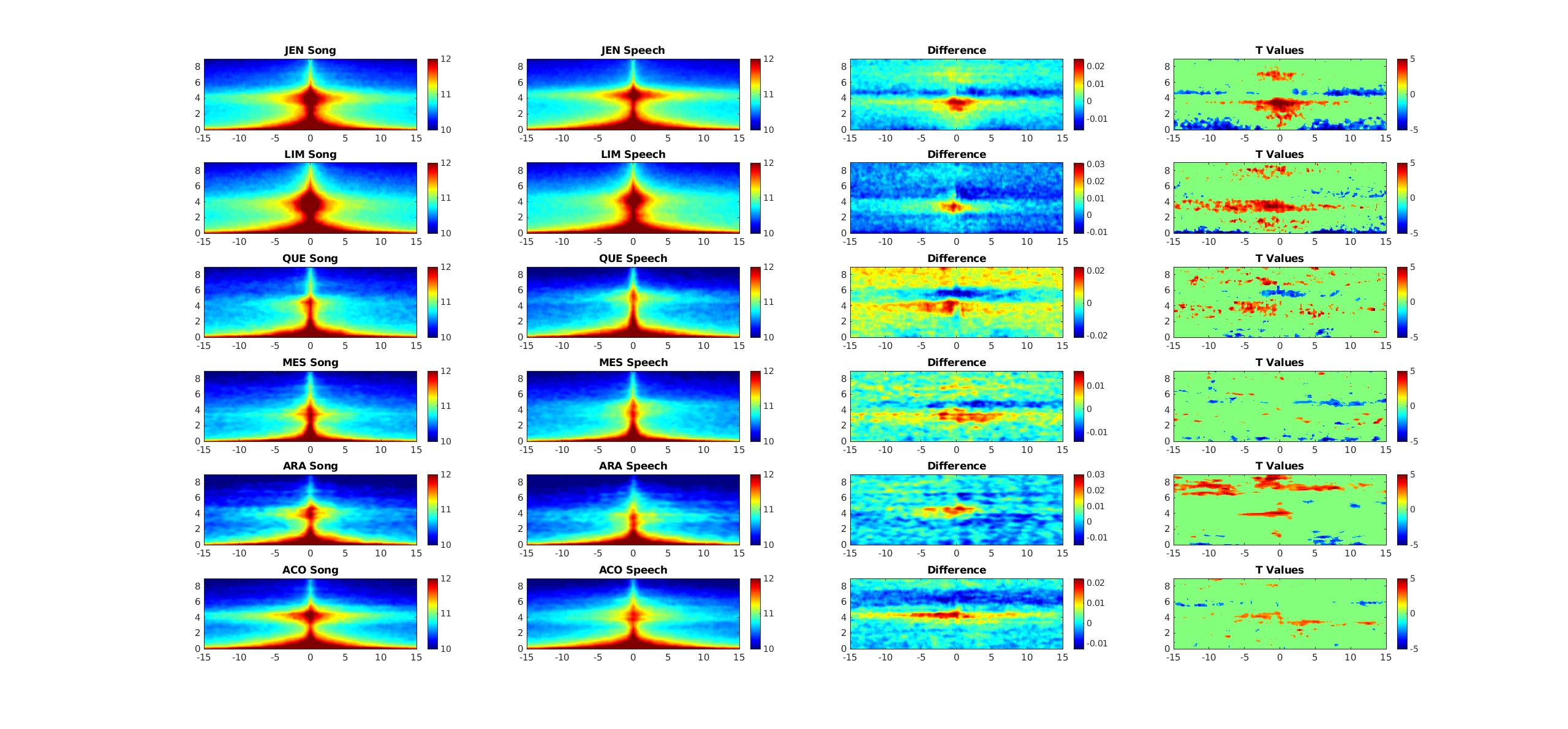

**Fig. S10. Extraction of spectro-temporal modulations patterns for singing and speaking vocalization samples.** Same as Fig. S7. Note that even for those societies in which differences did not reach statistical significance (mainly the last four), the trend was similar, as seen in the difference plots.

**Supplementary Discussion**

It is relevant to note that the phenomenon of lexical tone should be considered in the current study. Indeed, in languages with lexical tone, the discreteness of pitch variation in speech and song is more similar compared to languages without lexical tone. In the Hilton and al. database only two of the twenty-one societies included featured languages with lexical tone, and did not include languages with multiple level tones, such as Cantonese. In order to address this limitation we carried out both univariate and multivariate analyses on Cantonese song samples (20 samples recorded in Beijing) vs. Cantonese speech (20 sample from the Cantonese ASR database: <https://github.com/HLTCHKUST/cantonese-asr/tree/main/dataset> matched in gender and duration with the song samples, see Fig. S11 and S12 below).

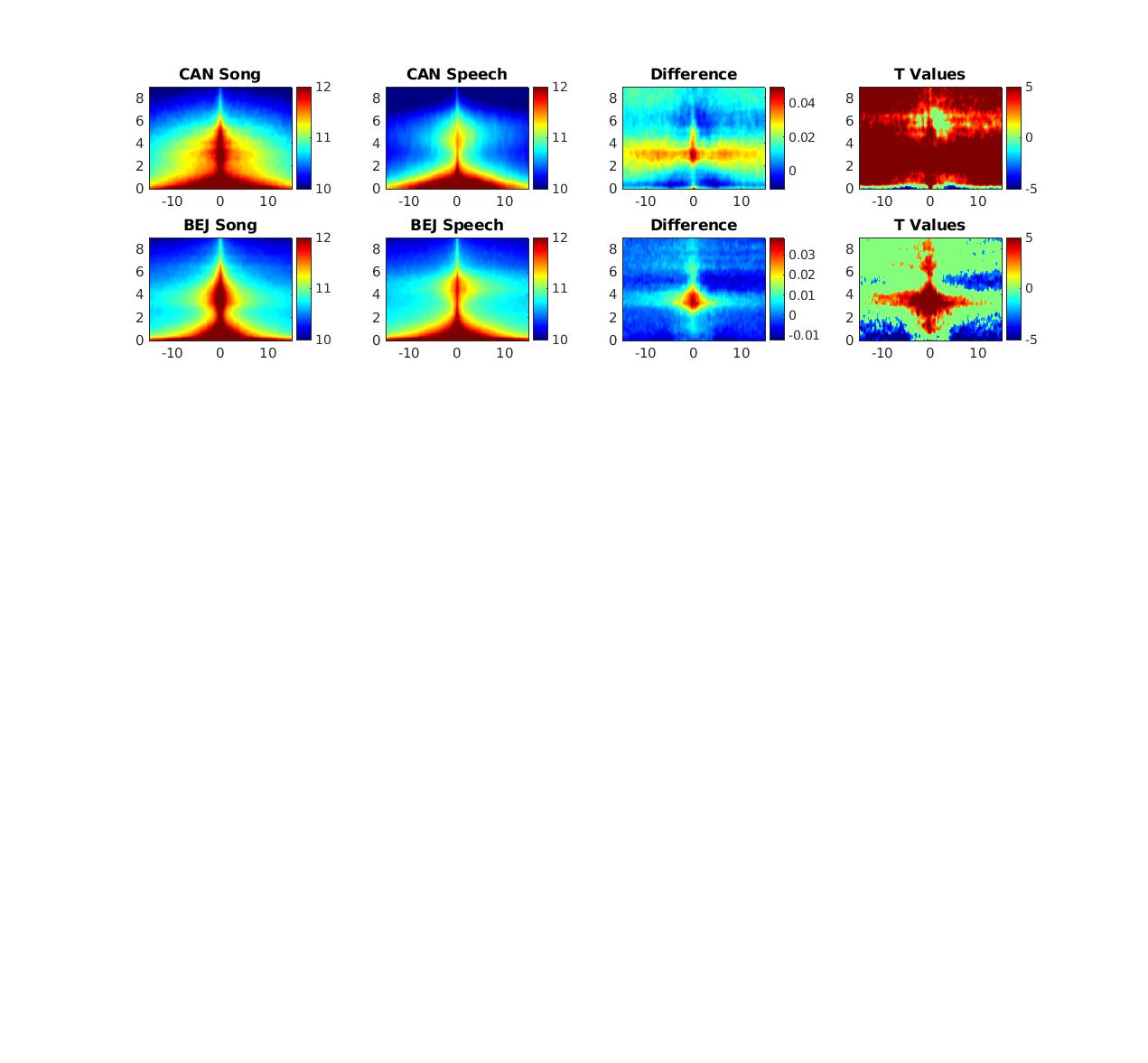

**Fig. S11.** Extraction of spectro-temporal modulations patterns for singing and speaking vocalization samples in the Cantonese sample**.** We extracted the spectrotemporal modulation patterns of the Cantonese sample and then averaged them. We computed the normalized difference between the song and speech modulation patterns. The panel in the third column therefore show where in the modulation space song differs from speech (red part of the scale) and where speech differs from song (blue part of the scale).

Univariate analysis (Song vs. Speech contrast of STM) revealed a clear overlap between the Cantonese analysis and the analysis performed with the 21 societies, within each category (song and speech) in the two specific acoustical ranges reported in the main text (see Fig. 2. and Figure S7 to S10). That is, the orange zones in the MPS, corresponding to energy found to be greater in song than speech largely fell within the range observed in the other 21 groups (black outlines from Fig. 2A). Similarly, the blue zones, corresponding to greater energy for speech compared to song, fall well within the range for speech in the univariate analysis from Fig 2A. So descriptively speaking, Cantonese is not different from what we expected.

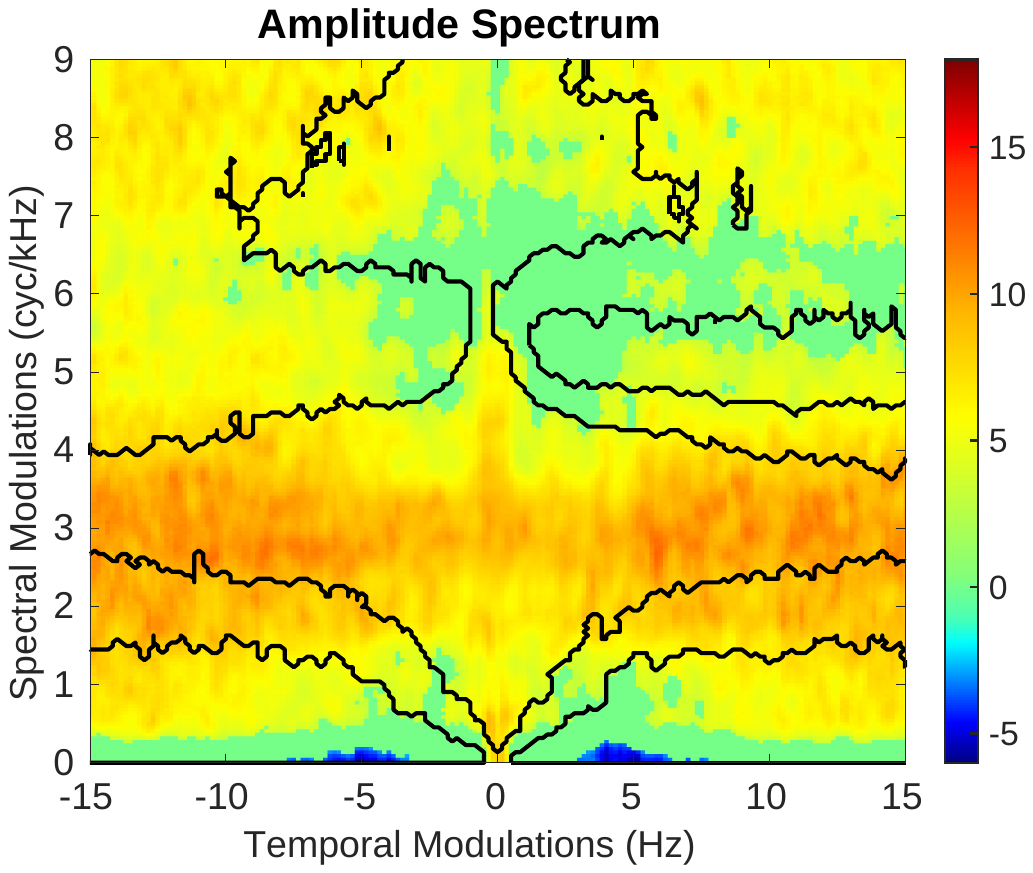

**Fig. S12.** Song vs. Speech contrast in the STM domain for Cantonese excerpts (p< .05, FDR-corrected). Dark lines illustrate the boundaries of the significant effects presented in Fig. 2A of the manuscript.

For multivariate results, we trained a classifier exclusively on the spectro-temporal features of the 21 societies reported in the manuscript (see methods) and then made predictions on Cantonese vocalizations. The model successfully classified Cantonese song and speech well above chance (accuracy = 80%; sensitivity = 85%, specificity = 75%). Thus, Cantonese speech and song share sufficient commonalities in terms of their spectrotemporal modulations with other societies that the model derived from those other societies generalizes quite well.

Overall, these results suggest the results reported in the main text can generalize even to a language with multiple lexical tones. There is one caveat, which is that we were unable to find a dataset containing Cantonese speech and song produced by the same speakers. From the samples we had available, we matched speech and song in terms of the gender of the speaker, but there are likely to be residual acoustical differences due to variable vocal characteristics. This effect, which we cannot easily quantify, would most likely add variability to our estimates of the STM of speech and music, and so the data we present above are probably a conservative estimate of the degree to which Cantonese is similar to the other groups. That is, if we had a nicely matched set of vocalizations, we expect the result would be even better than it is.

Table S1.

| Region | Sub-Region | Society | Language | Language family | Subsistence | Population | Distance to | Recordings |
| --- | --- | --- | --- | --- | --- | --- | --- | --- |
|  |  |  |  |  | type |  | city (km) |  |
| Africa | Central Africa | Mbendjele BaYaka | Mbendjele | Niger-Congo | Hunter- | 61-152 | 120 | 60 |
|  |  |  |  |  | Gatherer |  |  |  |
|  | Eastern Africa | Hadza | Hadza | Hadza | Hunter- | 35 | 80 | 38 |
|  |  |  |  |  | Gatherer |  |  |  |
|  |  | Nyangatom | Nyangatom | Nilotic | Pastoralist | 155 | 180 | 56 |
|  |  | Toposa | Toposa | Nilotic | Pastoralist | 250 | 180 | 60 |
| Asia | East Asia | Beijing | Mandarin | Sino-Tibetan | Urban | 21.5M | 0 | 124 |
|  | South Asia | Jenu Kurubas | Kannada | Dravidian | Other | 2000 | 15 | 80 |
|  | Southeast Asia | Mentawai | Mentawai | Austronesian | Horticulturalist | 260 | 120 | 60 |
|  |  | Islanders |  |  |  |  |  |  |
| Europe | Eastern Europe | Krakow | Polish | Indo-European | Urban | 771,069 | 0 | 44 |
|  |  | Rural Poland | Polish | Indo-European | Agriculturalists | 6,720 | 70 | 55 |
|  | Scandinavia | Turku | Finnish & Swedish | Uralic and | Urban | 186,000 | 0 | 80 |
|  |  |  |  | Indo-European |  |  |  |  |
| North | North America | San Diego | English (USA) | Indo-European | Urban | 3.3M | 0 | 116 |
| America |  |  |  |  |  |  |  |  |
|  |  | Toronto | English | Indo-European | Urban | 5.9M | 0 | 198 |
|  |  |  | (Canadian) |  |  |  |  |  |
| Oceania | Melanesia | Ni-Vanuatu | Bislama | Indo-European | Horticulturalist | 6,000 | 224 | 90 |
|  |  |  |  | Creole |  |  |  |  |
|  |  | Enga | Enga | Trans-New Guinea | Horticulturalist | 500 | 120 | 22 |
|  | Polynesia | Wellington | English (New | Indo-European | Urban | 210,400 | 0 | 228 |
|  |  |  | Zealand) |  |  |  |  |  |
| South | Amazonia | Arawak | English Creole | Indo-European | Other | 350 | 32 | 48 |
| America |  |  |  |  |  |  |  |  |
|  |  | Tsimane | Tsimane | Moseten-Tsimane | Horticulturalist | 150 | 234 | 51 |
|  |  | Sapara & Achuar | Quechua & Achuar | Quechuan & | Horticulturalist | 200 | 205 | 59 |
|  |  |  |  | Jivaroan |  |  |  |  |
|  | Central Andes | Quechua/Aymara | Spanish | Indo-European | Agro- | 200 | 8 | 49 |
|  |  |  |  |  | Pastoralist |  |  |  |
|  | Northwestern | Afrocolombians | Spanish | Indo-European | Horticulturalist | 300-1,000 | 100 | 53 |
|  | South America |  |  |  |  |  |  |  |
|  |  | Colombian | Spanish | Indo-European | Commercial | 470,000 | 0 | 43 |
|  |  | Mestizos |  |  | Economy |  |  |  |

**Table S1.**	Societies from which recordings were gathered, from ^6^, used with permission.

**Table S2**

| Label | Stub | Variables | Description | Significance |
| --- | --- | --- | --- | --- |
| Attack Curve Slope | mir_attack | Mean, Med, StD, Range, Min, Max, 1st Quart, 3rd Quart, IQR, Distance | MIRtoolbox detects acoustic events in the audio; for a subset of those it can compute an attack slope from amplitude curves, which is the slope of the line from the beginning of the event to its peak. | The slope of an attack curve provides a relative measure of "alerting components," or immediately discriminable beginnings of a vocalization. |
| Roughness | mir_roughness | Mean, Med, StD, Range, Max, 1st Quart, 3rd Quart, IQR, Distance | A roughness value produced by computing the peaks of the audio spectrum and taking the average of the dissonance between all possible pairs of peaks; following Buyens et al. (2017), we reduce this to a single measure by taking the  RMS-normalized mean. | Along with inharmonicity, roughness provides one measure of dissonance in a recording.  Roughness similarly provides at least one measure of vocal clarity. |
| 85th Energy Percentile | mir_rolloff85 | Whole | An estimate of the amount of high frequency in a signal measured by the frequency such that a 85% of the total energy is contained below it. | The 85th energy percentile allows a comparison of relative measures of high-frequency acoustics in a vocalization. |
| Inharmonicity | mir_inharmonicity | Whole | An estimate of the inharmonicity in the signal produced by identifying the number of partials that are not multiples of the fundamental frequency (i.e. those outside of the ideal harmonic range). | Along with roughness, inharmonicity provides a more precise measure of dissonance in a vocalization. |
| Tempo | mir_tempo | Whole | A tempo estimate made by detecting periodicities from MIR’s event detection curves. Outputs a single number. | Tempo allows assessment of the speed or pace of a vocalization. |
| Pule Clarity | mir_pulseclarity | Whole | Estimates the rhythmic clarity, or strength of the beats (Lartillot et al. 2008). | Pulse clarity provides a measure of the vocal clarity of a speaker or emphasis on individual utterances. |
| Rhythmic Variability | npvi_total | Recording | The nPVI equation measures the “average degree of durational contrast between adjacent events in a sequence" (Daniele & Patel, 2015). This makes it especially useful for comparing rhythmic units across language and music (i.e., syllables vs. notes). To automatically detect events, we used Mertens’ (2004) syllable detection algorithm. | By providing a measure of durational contrast, nPVI_total is a measure of rhythmic complexity in a recording. |
| Rhythmic Variability | npvi_phrase | Phrase | In addition to detecting syllables, Mertens’ algorithm detects phrases. Whereas npvi_total computes nPVI based on the whole file as a continuous phrase, this measure computes the nPVI for each detected phrase and reports the mean. In other words, it excludes the distances between the ends and beginnings of phrases. | nPVI_phrase provides a more granular measure of rhythmic complexity, within phrases, rather than between them. |

*(continued)*

| Label | Stub | Variables | Description | Significance |
| --- | --- | --- | --- | --- |
| Temporal Modulation | tm_peak_hz | Whole | The temporal modulation spectrum is the frequency decomposition of the amplitude envelope of a signal. This measures how loud something is at any given moment. We then measure how fast the loudness changes. For example: if someone sings a note every second, the spectrum will have a peak at 1Hz. If someone sings a note three times a second, but with an emphasis every three seconds, there will be a large peak at 1Hz, and a smaller peak at 3Hz. The peak of the spectrum is the frequency of the amplitude spectrum which has the highest root mean square of a given recording and represents a raw value of the recording’s tempo. | The peak of the temporal modulation spectrum provides a measure of how maximally modulated, or variable, the onset of notes are in a recording, providing a raw measure of metre for speech and song. |
| Temporal Modulation | tm_std_hz | StD | The temporal modulation spectrum is the frequency decomposition of the amplitude envelope of a signal. This measures how loud something is at any given moment. We then measure how fast the loudness changes. For example: if someone sings a note every second, the spectrum will have a peak at 1Hz. If someone sings a note three times a second, but with an emphasis every three seconds, there will be a large peak at 1Hz, and a smaller peak at 3Hz. The standard deviation of the spectrum is taken as a measure of how exaggerated the peak is. | The standard deviation of temporal modulation allows for an assessment of the overall variability of temporal modulations in a recording, providing a coarse measure of rhythm, with a lower standard deviation leaning towards more monorhythmic signals. |
| Pitch | praat_f0 | Mean, Med, StD, Range, Min, Max, 1st Quart, 3rd Quart, IQR | The fundamental frequency (f0) in Hertz for each recording | Pitch provides a fundamental measure of the highness or lowness, in frequency, of an utterance.  Likewise, the shape of the pitch curve and the overall value of pitch is a common discriminable feature  in both speech and song. |
| Pitch Space | praat_f0travel | Mean, Med, StD, Range, Max, 1st Quart, 3rd Quart, IQR | The distance between f0 at each  .03125/sec interval to the next. | Pitch space provides a dynamic measure of pitch’s range over time. |
| Pitch Rate | praat_pitch_rate | Whole, Med, IQR | The pitch rate is a measure of pitch change over time. In essence, the pitch rate provides a measure of pitch curve smoothness (a lower value corresponds to a smoother curve). | The pitch rate provides a measure of how smooth or variable pitch is over time. |
| Vowel Space | praat_vowtrav | Mean, Med, StD, Range, Max, 1st Quart, 3rd Quart, IQR | The Euclidian distance travelled in vowel space. This is equivalent to distance between the two formants. | Vowel space provides a measure of how much of the possible complex vowel space is used. |
| Vowel Space Travel Rate | praat_vowtrav_rate | Whole, Med, IQR | The Euclidian distance travelled in vowel space over a rate of time.  This is equivalent to distance  between two formants divided by rate of time. | Vowel travel rate provides a measure of how much of the vowel space is used over time, a relative measure of acoustic "flashiness" of a signal. |

*(continued)*

| Label | Stub | Variables | Description | Significance |
| --- | --- | --- | --- | --- |
| Amplitude | praat_intensity | Mean, Med, | A measure of amplitude (loudness) | Amplitude provides a measure of |
|  |  | StD, Range, | in decibels | how loud or quiet a vocalization is |
|  |  | Min, Max, 1st |  | and can be compared between |
|  |  | Quart, 3rd |  | types within speakers |
|  |  | Quart, IQR, |  |  |
|  |  | Distance |  |  |

Amplitude Space

Amplitude Rate

praat_intensitytravelMean, Med,

StD, Range, Max, 1st Quart, 3rd Quart, IQR

praat_intensity_rate Whole, Med,

IQR

The distance between amplitude at each .03125/sec interval to the next.

A measure of decay in intensity curves in each recording measured as change in amplitude over time.

Intensity space provides a dynamic measure of intensity’s range over time.

The intensity rate provides a measure of how loud or soft amplitude changes over time.

1st Formant	praat_f1	Mean, Med, StD, Range, Min, Max, 1st Quart, 3rd Quart, IQR

The frequency in Hertz of the 1st formant at each (.03125/sec) point

1st formants are the 1st in a harmonic series following from the fundamental frequency and is important for a number of acoustic reasons.

Second Formant

praat_f2	Mean, Med, StD, Range, Min, Max, 1st Quart, 3rd Quart, IQR

The frequency in Hertz of the second formant at each (.03125/sec) point

Second formants are the second in a harmonic series following from the fundamental frequency, and along with the 1st formant, is used by listeners to perceive vowels.

File duration	meta_length	The length of the unedited sound files

Concatenated file duration

meta_edit_length	The length of the concatenated versions of the sound files

Supplementary Table 2.	Codebook for acoustic features, from ^6^, used with permission
